# Single-cell omics-based characterization of human basophils reveals two transcriptionally distinct populations

**DOI:** 10.1101/2025.02.20.639059

**Authors:** Sofia Papavasileiou, Jiezhen Mo, Daryl Boey, Chenyan Wu, Magnus Tronstad, Remi-André Olsen, Jörg A. Bachmann, Jing Hui Low, Jocelyn Ong, Lars Heede Blom, Anand Kumar Andiappan, Gunnar Nilsson, Joakim S. Dahlin

## Abstract

**Background:** Basophils are implicated in various diseases including allergies, but a comprehensive characterization of human basophils has yet to be performed at the single-cell level.

**Objective:** We aimed to generate a single-cell omics-based resource of circulating human basophils, to be made accessible to the research community. We also sought to investigate basophil heterogeneity at the transcriptional and surface epitope levels.

**Methods:** Circulating basophils were analyzed using short- and long-read single-cell RNA-sequencing in combination with large-scale immunoprofiling of more than 100 cell surface markers.

**Results:** We used a cellular indexing of transcriptomes and epitopes by sequencing (CITE-seq) based cell barcoding system to accurately track side scatter^low^ lineage^−^ CCR3^+^ Fc_ε_RI^+^ basophils through the single-cell omics analysis. The approach allowed for accurate transcriptional profiling of basophils among low-density peripheral blood leukocytes. The generation of an additional dataset with only basophils revealed two transcriptionally distinct populations. Single-cell transcriptomic analysis of samples from additional donors was performed and coupled with a large-scale immunoprofiling screen, producing a third CITE-seq dataset containing basophils. The analysis verified the existence of the two transcriptionally distinct populations and revealed that these populations have similar immunophenotypes with regard to the investigated surface markers. Long-read single-cell RNA-sequencing analysis provided further insights into the gene expression dynamics of the circulating basophils and confirmed the transcriptionally defined basophil states.

**Conclusions:** The single-cell omics analysis revealed two transcriptionally distinct basophil populations. Our single-cell omics resource constitutes a reference for charting human basophils at the cellular and molecular levels in health and disease.

**Capsule summary:** Short- and long-read single-cell RNA-sequencing coupled with large-scale immunoprofiling characterize human basophils in peripheral blood and reveal two transcriptionally distinct populations.

## Introduction

Single-cell RNA-sequencing technologies allow for profiling of thousands of individual cells in parallel. Large-scale application of these technologies has allowed consortia, such as the Human Cell Atlas,^1^ to establish reference collections of transcriptomics data of millions of cells. These collections enable the wider research community to gain cellular and molecular insights into the cell types throughout the body. Virtually all broad classes of immune cells have been characterized at the single-cell transcriptional level, including the allergy-associated eosinophil^2–4^ and mast cell^5, 6^ populations. However, a detailed characterization of basophils has yet to be performed at the single-cell transcriptional level.

Here, we perform a comprehensive single-cell omics-based analysis of human basophils in peripheral blood. We show that cellular indexing of transcriptomes and epitopes by sequencing^7^ (CITE-seq) analysis detects basophils among circulating leukocytes. Single-cell RNA-sequencing coupled with large-scale immunoprofiling reveals two transcriptionally distinct basophil states and shows that the identified populations harbor similar immunophenotypes. Long-read-based single-cell RNA-sequencing verifies the transcriptionally distinct populations using an independent sequencing approach and provides further insights into the basophils’ gene expression dynamics. The processed single-cell omics data is made accessible through the Gene Expression Omnibus, GSE287976, and through a user-friendly web interface (http://dahlinlab.cmm.se). A detailed methods section is provided in supplement.

## Results and Discussion

The capacity of the single-cell RNA-sequencing workflow to detect the human basophil population has yet to be evaluated and strategies to confidently annotate the cell population in datasets are missing. We therefore devised a CITE-seq-based strategy to track primary basophils from peripheral blood throughout the experiment, from the input sample to the resulting dataset. Briefly, we sorted fluorescently and oligonucleotide-barcoded basophils (side scatter^low^ lineage^−^ CCR3^+^ Fc_ε_RI^+^ cells^8^) and mixed them with low-density leukocytes (Fig 1, A). Flow cytometry analysis showed that the cell suspension used as input for the single-cell RNA-sequencing contained 26 % barcoded basophils (Fig 1, B). As comparison, 17 % of the cells in the single-cell dataset constituted barcoded cells after quality control and filtering (Fig 1, C, Fig E1), revealing successful capture of the basophil population with a small selective loss of these cells.

**FIG 1.**
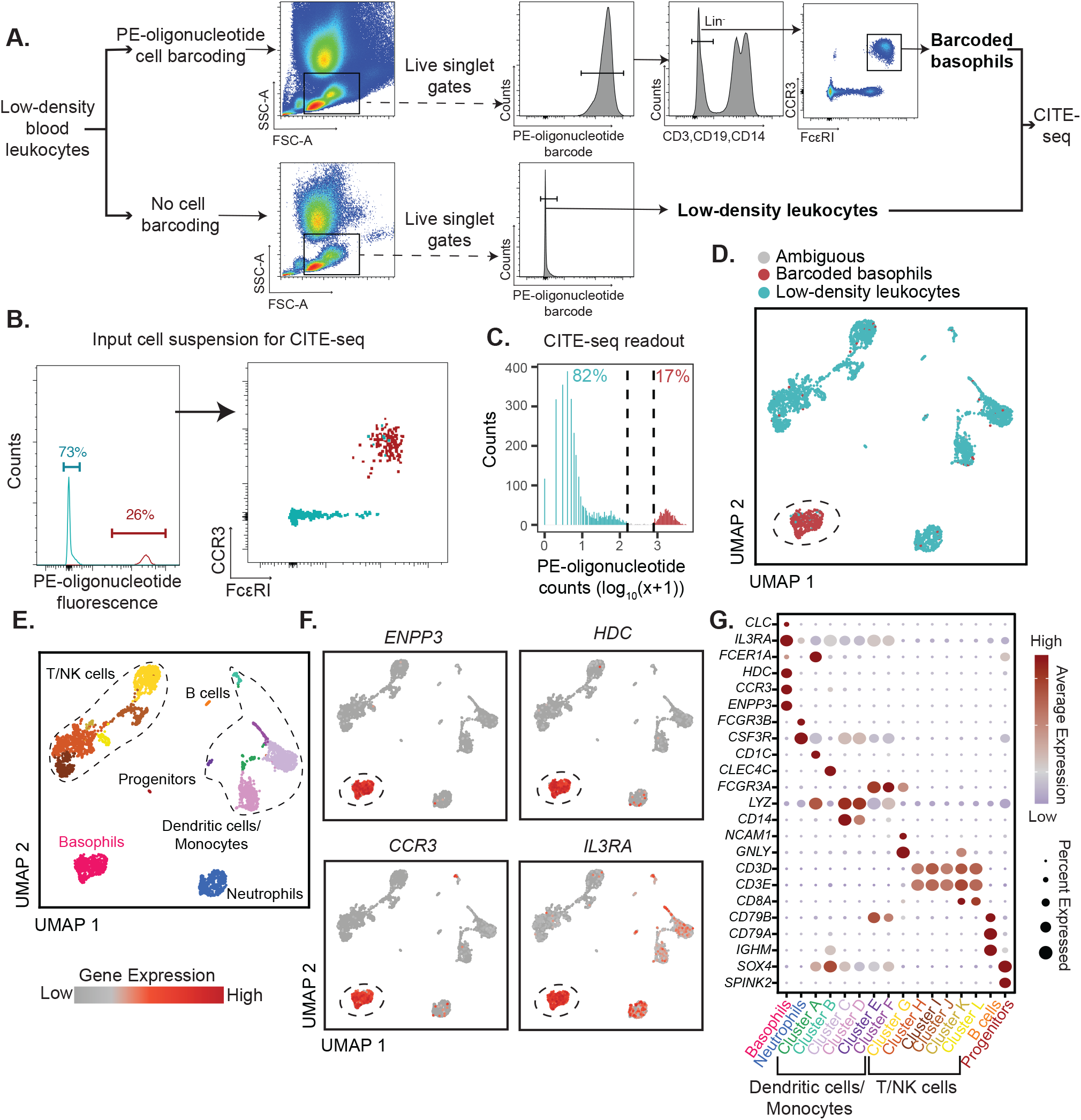
Single-cell RNA-sequencing captures the gene expression profile of circulating basophils. (**A**) Experimental design. (**B**) Flow cytometry analysis showing the proportion of barcoded cells prior to CITE-seq analysis. The plots show live cells. (**C**) Analysis of CITE-seq data showing the proportion of barcoded cells. The histogram shows the distribution of the PE-oligonucleotide derived sequencing reads after the quality control and filtering steps. (**D**) UMAP visualization of the single-cell transcriptomics dataset (3720 cells), colored according to panel C. (**E**) Cells colored according to Leiden clusters at resolution 0.4. The clusters are annotated based on marker genes in panel G. (**F**) Normalized gene expression of selected genes. The basophil cluster is encircled. (**G**) Dot plot generated with the DotPlot function of the Seurat package showing the scaled average expression of cell-type related genes of the clusters shown in panel E. The colors of the x-axis labels correspond to the colors of the Leiden clusters shown in panel E.

We visualized the single-cell transcriptomics data using UMAP plots (Fig 1, D-F). The absence of a single-cell-based reference to annotate the basophil population based on gene expression profile prompted us to specifically plot the oligonucleotide-barcoded cells (Fig 1, D). These cells constituted the flow cytometry-defined basophils, allowing for accurate identification of the basophil cluster (Fig 1, D-E). Plotting the expression levels of cell type-related genes annotated the remaining clusters, which included lymphocyte populations, dendritic cells/monocytes, and neutrophils (Fig 1, E-G). As expected, the basophil cluster expressed genes coding for basophil-related markers CD203c (*ENPP3*), histidine decarboxylase (*HDC*), CCR3 (*CCR3*), and the IL-3 receptor (*IL3RA*) (Fig 1,F). Taken together, the results revealed the successful analysis of basophils using single-cell RNA-sequencing.

We generated an additional dataset comprising basophils as the only cell type to investigate the transcriptional heterogeneity of basophils. The flow cytometry analysis, performed as part of the basophil isolation procedure, revealed a gradient of c-Kit expression on the cells (Fig 2, A). We therefore sorted and mixed the entire basophil population (side scatter^low^ lineage^−^ CCR3^+^ Fc_ε_RI^+^ cells) with oligonucleotide-barcoded c-Kit^+^ basophils for the CITE-seq analysis (Fig 2, A, Fig E2, A). Strikingly, two transcriptionally distinct clusters of basophils were identified following data processing and visualization (Fig 2, B). These populations were referred to as Basophils 1 and 2. The c-Kit^+^ basophils were dispersed among Basophils 1 and 2, indicating that high c-Kit expression is uncoupled from transcriptional heterogeneity (Fig 2, C, Fig E2). Differential gene expression analysis revealed higher levels of *CLC* and *FTL* in Basophils 2 than in Basophils 1, whereas the expression levels of the basophil-associated genes *CCR3* and *HDC* were comparable between the clusters (Fig 2, D-E). As an initial test to validate the presence of two populations, we mapped cells from the basophil cluster in the first dataset onto the present second dataset (Fig 2, F). The results revealed that the basophils from the first dataset mapped onto both Basophils 1 and 2, with patterns of high *CLC* and *FTL* expression in the Basophil 2 cluster, supporting the observation of two transcriptionally distinct basophil populations (Fig 2, G-H).

**FIG 2.**
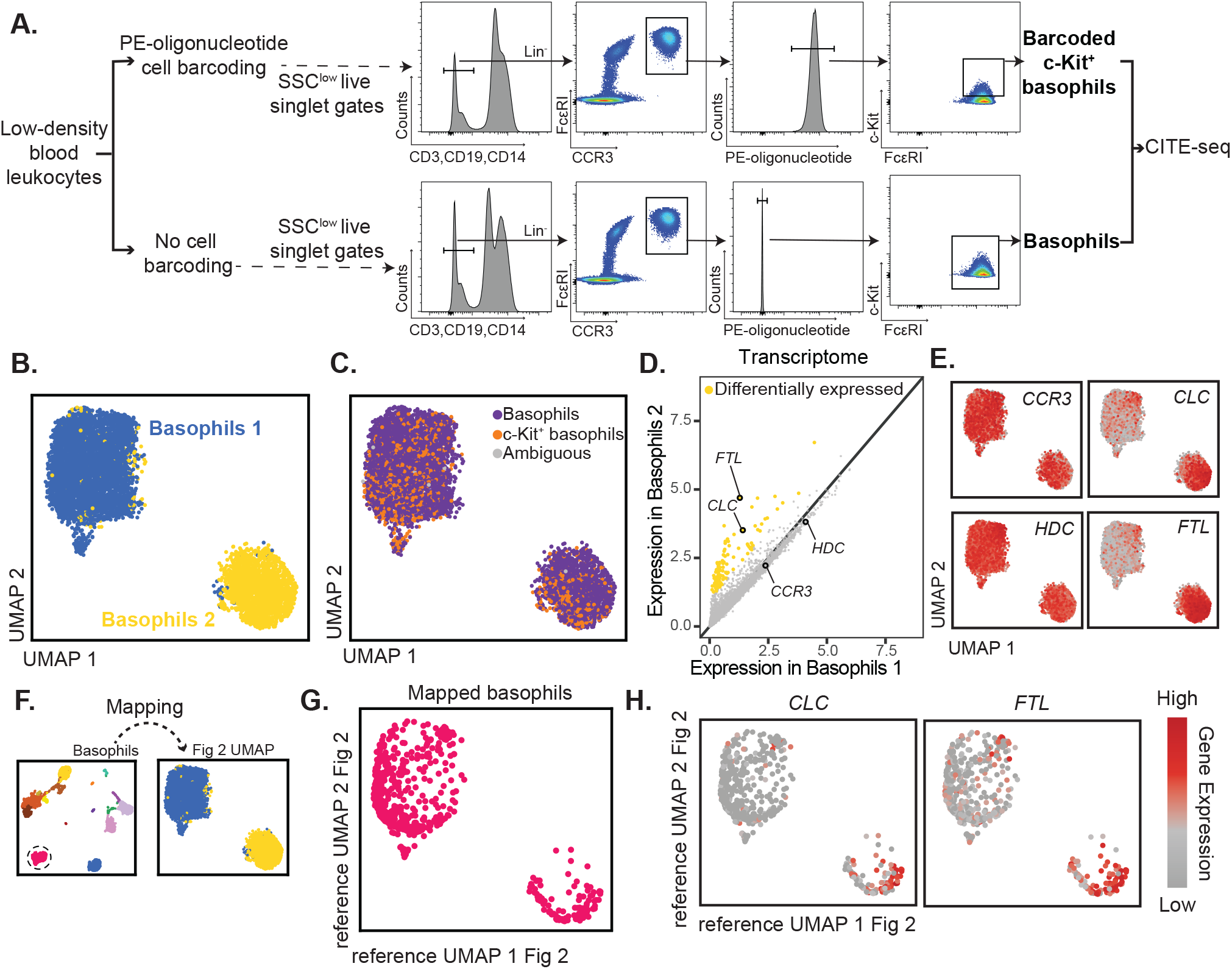
Circulating basophils constitute two transcriptionally distinct populations. (**A**) Experimental design. (**B**) UMAP visualization of the 6285 basophils with the colors representing Leiden clusters calculated at a resolution 0.1. (**C**) Cells colored by the sorting strategy in panel A and Fig E2. **(D)** Scatter plot showing the mean normalized gene expression of the two basophil populations on a log_2_(x+1) scale according to the AverageExpression function. The colored dots represent differentially expressed genes. (**E**) UMAP visualizations showing the normalized gene expression. (**F**) Schematic representation of the mapping of the annotated basophils from Fig 1 onto the UMAP plot of panel B. (**G**) UMAP visualization showing the basophils from Fig 1 mapped onto the UMAP space of the sorted basophils in Fig 2. (**H**) Gene expression levels of the mapped basophils.

The unexpected discovery of two transcriptionally distinct basophil populations warranted further investigation and the generation of an additional dataset from independent donors. To assess the robustness of observing the two populations, we changed the cell isolation procedure from fluorescence-activated cell sorting-based to magnetic bead-based basophil purification. A CCR3-based isolation method was chosen, as it allows the isolation of the entire basophil population, and density gradient centrifugation ensured that virtually no CCR3^+^ eosinophils remained before the single-cell sequencing protocol (Fig E3, A-B). The CCR3-enriched basophils were incubated with a panel of 140 different oligonucleotide-conjugated antibodies prior to the CITE-seq analysis to maximize the insights gained from the additional samples (Fig 3, A). Three replicates from two independent donors together formed a comprehensive multimodal dataset of basophils, with transcriptomics data coupled with large-scale immunoprofiling data of each single cell.

**FIG 3.**
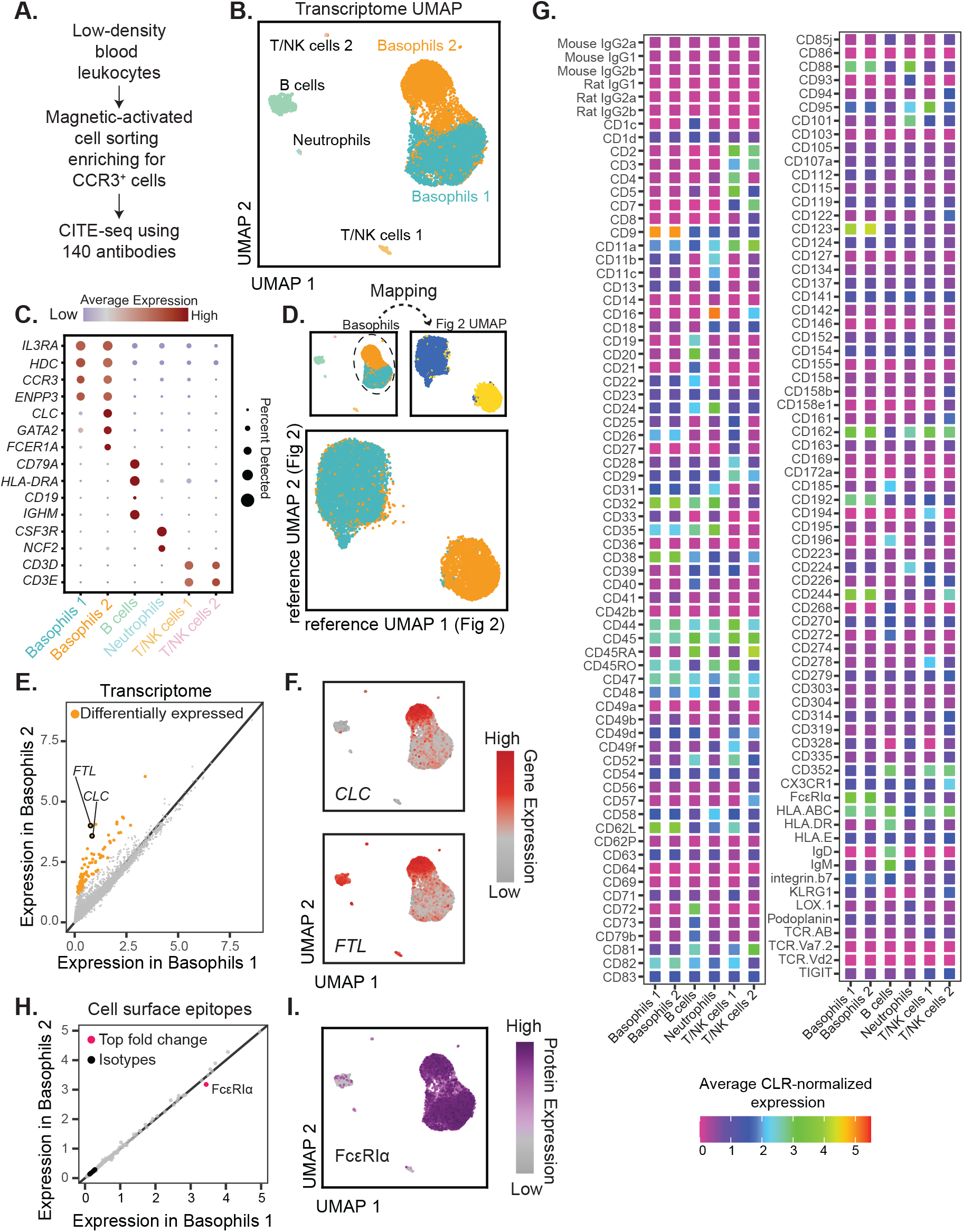
A multimodal resource reveals two transcriptionally distinct basophil populations with similar immunoprofiles. (**A**) Experimental design. (**B**) UMAP visualization of 15795 cells from 2 donors and 3 CITE-seq libraries. The colors indicate annotated Leiden clusters at resolution 0.1. (**C**) Dot plot generated with the DotPlot function of the Seurat package showing the scaled average gene expression of cell-type-defining genes, used to annotate the cells in panel B. (**D**) UMAP visualization of the basophils from panel B mapped onto the UMAP space of the sorted basophil dataset of Figure 2. The colors correspond to the Leiden clusters shown in panel B. (**E**) Scatter plot showing the mean normalized gene expression of the two basophil populations, plotted on a log_2_(x+1)-scale according to the AverageExpression function. The colored dots represent differentially expressed genes. **(F)** UMAP visualizations of the normalized gene expression. **(G)** Heatmap showing the average CLR-normalized expression of 140 oligonucleotide-conjugated antibodies. (**H**) Scatter plot showing the mean of the CLR-normalized cell surface epitope expression of the two basophil populations. **(I)** UMAP visualization of the CLR-normalized protein expression of Fc_ε_RIα, the protein with the highest fold change.

Transcriptome-based UMAP visualization and cluster annotation revealed two basophil populations in the integrated multimodal dataset (Fig 3, B-C), similar to the reference dataset (Fig 2, B). Notably, the observed basophil heterogeneity was independent of replicate- or cell cycle-driven effects (Fig E3, C-D). The presence of two basophil clusters was replicated within each of the three constituent datasets when they were processed separately (Fig E3, E-F). Mapping the basophils from the integrated multimodal dataset onto the reference (Fig 2) showed matching Basophil 1 and 2 clusters across datasets (Fig 3, D), in which high *FTL* and *CLC* expression marked the Basophil 2 cluster (Fig 3, E-F). These observations confirmed the existence of the two transcriptionally distinct basophil populations in multiple donors and datasets and using different cell isolation methods.

Vivanco Gonzales et al performed a mass cytometry-based characterization of peripheral blood basophils, simultaneously measuring the expression of 44 proteins.^9^ The study identified 4 potential basophil states, distinguished based on the expression levels of CD16, CD244, and FcεRI. Our CITE-seq-based approach characterized in excess of 100 cell surface epitopes, including the above-mentioned markers. However, the expression levels of all markers were comparable between Basophils 1 and 2 (Fig 3, G-I). This included markers that are associated with basophil activation, such as CD11b, CD44, CD54, CD63, and CD69.^10^ We plotted the expression of surface epitopes to investigate whether any marker distinguished basophil subsets irrespective of cluster attribution. FcεRI constituted 1 out of 4 markers that exhibited significant heterogeneity (Fig E4), a marker also reported by Vivanco Gonzales et al.^9^ However, the FcεRI expression in basophils correlates with serum IgE levels^11^ and further inspection revealed that donor variability caused the bimodal FcεRI expression pattern (Fig E4). Notably, only CD352 subsets the basophils on the surface epitope level irrespective of donor (Fig E4, B). Flow cytometry analysis of CD352 verifies this observation.^12^ However, the CD352 levels were comparable in the Basophils 1 and 2 clusters (Fig 3, G).

Single-cell RNA-sequencing based on the Oxford Nanopore Technology (ONT) has the potential to capture full-length transcripts. We reasoned that this long-read sequencing technology could potentially improve the detection of specific genes, thereby gaining insights into basophil heterogeneity beyond the results obtained from the default short-read Illumina-based sequencing pipeline. Results based on the Nanopore platform would also enable us to validate the existence of basophil populations using an independent sequencing technology. To generate the long-read sequencing data, we produced Nanopore sequencing libraries from the existing PCR-amplified full-length cDNA libraries used for generating the results presented in Fig 3 – one from each donor. The preservation of the cell barcodes on the cDNA sequences allowed comparative analysis between the Nanopore- and Illumina-based results (Fig 4, A, Fig E5). UMAP visualization of the Nanopore data revealed three main basophil clusters (Fig 4, B). Two Nanopore clusters, referred to as Basophils 1 and Basophils 1 ONT, mainly comprised cells annotated as Basophils 1 according to the Illumina-based analysis (Fig 4, C). The discrimination of the Nanopore-specific Basophil 1 ONT cluster was attributed to two long non-coding RNAs (Fig 4, D). The Nanopore Basophils 2 cluster mainly comprised cells of the corresponding Illumina cluster (Fig 4, B-C, Fig E5). Differential expression analysis based on the Nanopore data revealed higher expression of *CLC* and *FTL* in Basophils 2 than in Basophils 1 (Fig 4, E-F), in agreement with the Illumina-based results (Fig 2, D-E, Fig 3, E-F). We also detected higher levels of *TSPAN9* and *OOEP* in Basophils 2 than in Basophils 1 in the Nanopore dataset, genes that were poorly captured in the Illumina dataset despite the overall higher sequencing depth (Fig 4, G-H, Table E1). These observations highlight the advantage of using two independent sequencing pipelines.

**FIG 4.**
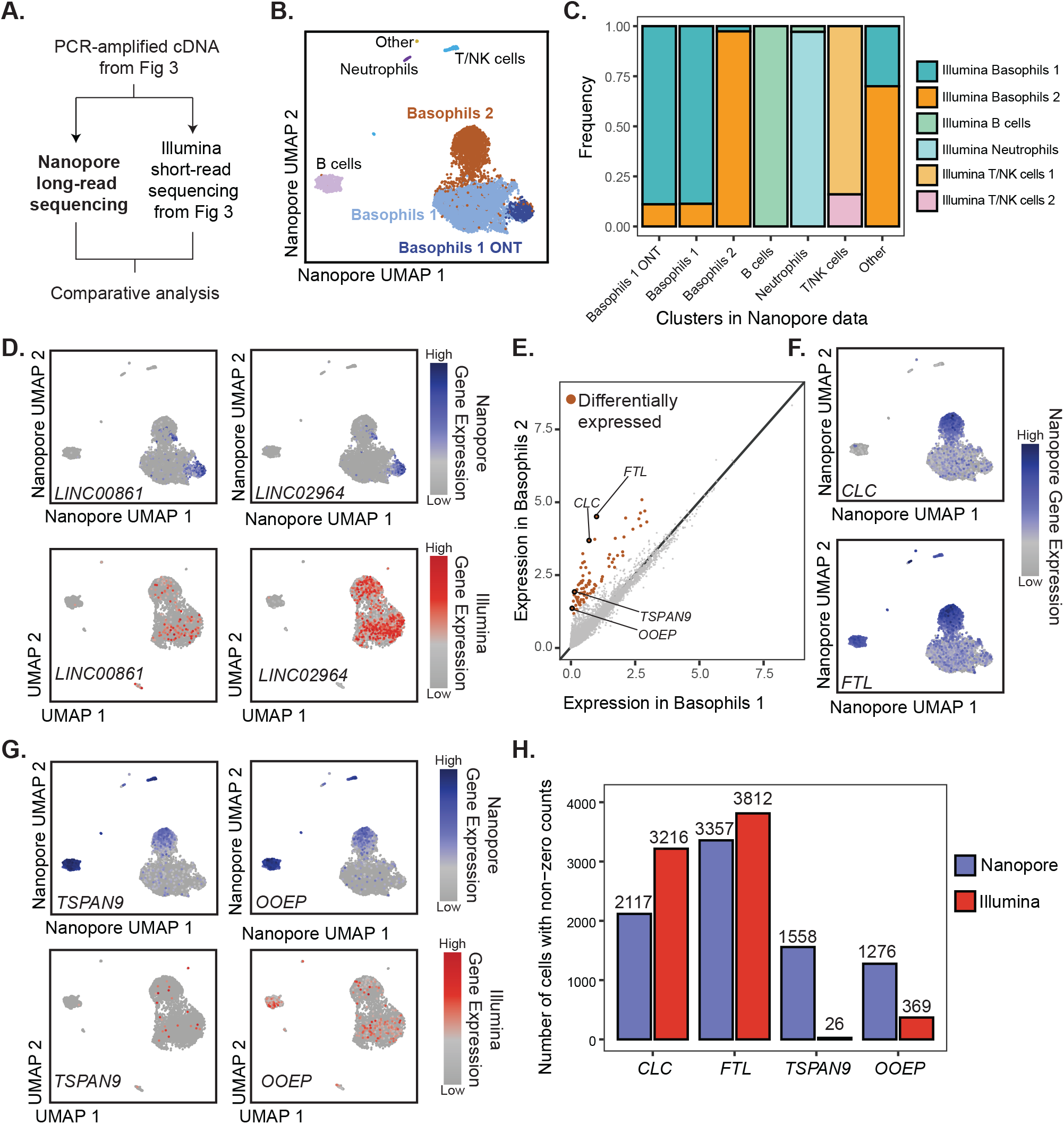
Long-read single-cell RNA sequencing provides additional insights into the basophils’ transcriptome. **(A)** Experimental design. **(B)** UMAP visualization of the long-read sequencing data showing 7304 cells from 2 donors. The colors indicate annotated Leiden clusters at resolution 0.1. **(C)** Bar plot showing the cell composition of the Nanopore-defined clusters (x-axis) according to the Illumina cluster annotation. **(D)** UMAP visualizations showing the normalized gene expression based on Nanopore and Illumina sequencing data. Note that only the cells that were profiled with both sequencing pipelines (7304 cells) are shown in the plots. **(E)** Scatter plot showing the Nanopore-based mean normalized gene expression in the two basophil populations on log_2_(x+1) scale according to the AverageExpression function. The colored dots highlight differentially expressed genes. **(F-G)** UMAP visualization showing normalized gene expression based on Nanopore and Illumina sequencing data. Only the cells profiled by both technologies are shown. **(H)** Bar plot showing the number of cells with at least one sequencing read, plotted separately for each sequencing pipeline. Only the cells profiled with both technologies were interrogated.

To conclude, single-cell short- and long-read-based RNA-sequencing coupled with large-scale immunoprofiling detects two transcriptionally distinct basophil populations. The single-cell omics data presented here constitutes a resource for future studies on basophils in health and disease.

## Acknowledgements

The research was funded by grants from the Swedish Research Council (2022-00558), the Swedish Cancer Society, and Karolinska Institutet. S.P. was supported by a scholarship from the Onassis Foundation (scholarship ID: F ZO 057-2/2022-2023). A.K.A., J.H.L., and J.O. were supported by core funds from the Singapore Immunology Network, Agency for Science, Technology and Research (A*STAR). The authors acknowledge support from the National Genomics Infrastructure in Stockholm funded by Science for Life Laboratory, the Knut and Alice Wallenberg Foundation and the Swedish Research Council, and SNIC/NAISS/Uppsala Multidisciplinary Center for Advanced Computational Science for assistance with massively parallel sequencing and access to the UPPMAX computational infrastructure. Sequencing data was processed using resources provided by NAISS, partially funded by the Swedish Research Council through grant agreement no. 2022-06725.

