## Supplement for "Single-cell omics-based characterization of human basophils reveals two transcriptionally distinct populations"

### Methods

*Study design*

The study was designed to analyze basophils at a progressively deeper level in each single-cell omic dataset, allowing for validation of key findings across datasets. Successful CITE-seq analysis of basophils among low-density leukocytes is shown in Figure 1 – results from 1 independent buffy coat. CITE-seq analysis of basophils as the only cell type is presented in Figure 2 – results from 1 independent buffy coat. CITE-seq analysis including a large-scale immunoprofiling of basophils is shown in Figure 3 – results from 3 single-cell libraries based on 2 independent buffy coats. Two of the PCR-amplified full-length cDNA libraries used to generate the data presented in Figure 3 were repurposed to generate the Nanopore sequencing dataset (one library for each buffy coat). This strategy allowed for a direct comparison between the short-read Illumina and the long-read Nanopore pipelines (Table E1).

*Ethics statement*

The Swedish Ethical Review Authority approved the study, and the subjects provided informed consent before sample collection.

*Sample processing and cell isolation*

The samples were from subjects that fulfilled the requirements to be blood donors in Sweden, which includes criteria of being at least 18 years old, weighing at least 50 kg, and being healthy. The buffy coat fraction of peripheral blood from the donors was processed. Low-density leukocytes were isolated using Ficoll-Paque density gradient centrifugation. Any remaining red blood cells were removed using BD Pharm Lyse (BD Biosciences, Franklin Lakes, New Jersey, USA).

*Flow cytometry*

Cells were incubated with antibodies against CD3 (clone HIT3a), CD19 (HIB19), CD14 (M5E2), CCR3 (5E8), FcεRI (CRA-1), CD117 (A3C6E2). 4',6'-diamidino-2-phenylindole (DAPI) or 7-aminoactinomycin D (7-AAD) was used to exclude dead cells. The fluorescently labelled antibodies and live/dead markers were purchased from Biolegend (San Diego California, USA), BD Biosciences (Franklin Lakes, New Jersey, USA), and Miltenyi Biotec (Bergisch Gladbach, Germany). The flow cytometry data was analyzed using FlowJo software (Version 10.10.0, BD Life Sciences, Ashland, Oregon, USA).

*Generation of the dataset presented in Figure 1*

Low-density leukocytes were stained with fluorescently labelled antibodies against CD3 (HIT3a), CD19 (HIB19), CD14 (M5E2), CCR3 (5E8), FcεRI (CRA-1), and the oligonucleotide-labeled antibodies TotalSeq-B0391 anti-human CD45 and TotalSeq-B0090 Mouse IgG1 κ isotype control (Biolegend). The stained cells were split into two cell fractions. One cell fraction was stained with biotinylated β2-microglobulin antibodies (2M2) and the other fraction with unconjugated β2-microglobulin antibodies (2M2) (Biolegend). Each cell fraction was then incubated with TotalSeq-B0953 PE streptavidin (Biolegend). Antibody incubation steps prior to the fluorescence-activated cell sorting (FACS) were performed in the presence of 2 mM EDTA. We sorted side scatter$^{low}$ (SSC$^{low}$) Lin$^{-}$ PE$^{+}$ CCR3$^{+}$ FcεRI$^{+}$ basophils from the PE-labeled cell fraction (Fig 1, A top) and SSC$^{low}$ PE$^{-}$ leukocytes from the unlabeled cell fraction (Fig 1, A bottom) using the FACS Aria Fusion system (BD Biosciences). The two cell fractions were pooled and CITE-seq analysis was performed. The cells were maintained at 4 °C or on ice after the isolation of low-density leukocytes.

*Generation of the dataset presented in Figure 2*

Low-density leukocytes were split into two fractions. Each cell fraction was incubated with either biotinylated or unconjugated β2-microglobulin antibody (2M2), followed by the addition of TotalSeq-B0953 PE streptavidin together with the fluorescently labelled antibodies against CD117 (A3C6E2), CD3 (HIT3a), CD19 (HIB19), CD14 (M5E2), CCR3 (5E8), and FcεRI (CRA-1). Antibody incubation steps prior to FACS were performed in the presence of 2 mM EDTA. Unlabeled basophils ($Lin^-$ $PE^-$ $CCR3^+$ $FcεRI^+$) and $PE^+$ $c\text{-}KIT^+$ basophils ($Lin^-$ $PE^+$ $CCR3^+$ $FcεRI^+$ $c\text{-}Kit^+$) were sorted using the FACS Aria Fusion system (BD Biosciences). The sorted cells were pooled and stained with oligonucleotide-labeled antibodies that included TotalSeq-B0090 Mouse IgG1 isotype control and TotalSeq-B0391 anti-human CD45 before the CITE-seq analysis (Biolegend). The cells were maintained at 4 °C or on ice after the isolation of low-density leukocytes.

*Generation of the dataset presented in Figure 3*

Low-density leukocytes were incubated with CCR3-APC-Cy7 antibodies (Biolegend) followed by staining with anti-Cy7 magnetic beads (Miltenyi Biotec). Magnetic-activated cell sorting enriched the $CCR3^+$ cells. The cells were maintained at 4 °C during the staining and magnetic separation. An additional Ficoll density gradient centrifugation was performed at room temperature to remove dead cells. The $CCR3^+$ cells were then maintained at 4 °C or on ice during the remaining steps. We incubated the cells with the TotalSeq-B Human Universal Cocktail v1.0 (Biolegend) according to the manufacturer's recommendations. The $CCR3^+$ cells were further enriched and washed using a magnetic column. Flow cytometry analysis using the BD FACSCanto II system (BD Biosciences) was performed on a fraction of the cells to verify cell purity. The antibody incubation and magnetic separation steps were performed in the presence of 2 mM EDTA. However, the EDTA was removed before the CITE-seq workflow as it interferes with the cDNA synthesis.

*Short-read single-cell transcriptomics data generation*

Single-cell datasets were generated with the Chromium Next GEM Single Cell 3' Reagent Kit v3.1 (dual index) with Feature Barcode technology for Cell Surface Protein (10X Genomics, Pleasanton, California, USA). To ensure the capture of the basophils, known to have low RNA contents,[1] one additional PCR cycle was introduced in the first cDNA amplification step. The final libraries were sequenced on the Novaseq 6000 platform (Illumina, San Diego, California, USA).

*Bioinformatic analysis*

The fastq files were processed with Cell Ranger (10X Genomics). The data in Figure 2 was processed with Cell Ranger version 5.0.1 with the option --include-introns. The datasets in Figures 1 and 3 were processed with Cell Ranger version 7.0.1, in which intron-containing reads are included by default. The reads were mapped against the GRCh38 reference genome (refdata-gex-GRCh38-2020-A).

The Cell Ranger-derived raw count tables with all the detected cell barcodes were used for the data analysis. Briefly, the count tables were imported into R (version 4.3.3) and analyzed with Seurat (version 4.3.0.1).[2] The analysis was performed using RStudio (version 2024.12.0+467). A recent study detected cells with low gene expression levels using antibody reads.[3] Inspired by this approach, cell barcodes were filtered based on the raw reads of the CD45 surface marker expression, a marker expressed on all leukocytes (Table E2). To identify barcoded and non-barcoded cells in Figures 1 and 2, we used the (raw) number of TotalSeq-B0953 PE streptavidin antibody reads (Table E2).

Cell barcodes with low numbers of detected features were removed and cell doublets were simulated using scDblFinder[4] (Version 1.10.0) with default parameters. The cell barcodes that were predicted to constitute doublets were removed from the analysis. Additional quality control filtering steps included the removal of cells showing high numbers of detected features and/or high percentage of mitochondrial transcript reads, indicative of dead or dying cells. Quality control and doublet removal steps were performed independently for each dataset to ensure strict thresholds appropriate for each sample (Table E2). Genes detected in less than 5 cells were removed (related to Fig 1-2). To generate the integrated dataset (related to Fig 3), genes detected in less than 30 cells were removed in the combined object.

The gene expression counts were normalized using the NormalizeData function with default parameters, and variable features were selected using the vst method. The number of leading principal components that were used in each dataset was determined by using the Jackstraw method.[5] To integrate the datasets in Figures 3 and 4 respectively, the Harmony[6] package (version 0.1.1) was used. The Leiden algorithm was used for clustering, with resolutions described in each Figure legend. The data was visualized using Uniform Manifold Approximation and Projection (UMAP),[7] in which the number of nearest neighbors and principal components were selected to match the values used for clustering. Surface epitope data was centered log ratio (CLR) normalized using the NormalizeData function.

The differential expression analysis between the two basophil populations was performed using the *FindMarkers* function using default parameters (i.e. normalized counts) and specifying the two groups to be compared. Genes and surface markers were considered differentially expressed if the adjusted p-values were smaller than or equal to 0.01 and the average log2 fold change values were larger than or equal to 1.

The cell cycle phase attribution was performed using the *CellCycleScoring* function and the Seurat-associated gene list (cc.genes.updated.2019).

To compare cells between datasets, we mapped the data from Figures 1 and 3 into the dataset of Figure 2. The dataset of Figure 2 was set as reference and the datasets of Figures 1 and 3 were set as queries. The PCA calculated prior to Harmony processing step was used to map the dataset from Figure 3. Integration anchors were calculated using the function *FindTransferAnchors* with default parameters. The query datasets were mapped onto the reference UMAP using the calculated anchors with the function *MapQuery* including the argument *reduction.model=”umap”*.

*Long-read sequencing*

PCR-amplified full-length cDNA remaining from the CITE-seq analysis of Figure 3 was used to generate the long-read sequencing libraries. To obtain sufficient material for the Nanopore Library preparation we further amplified the full-length cDNA using the cDNA primers (PN 2000089), provided with the Chromium Next GEM Single Cell 3’ Reagent Kit v3.1 (dual index) with Feature Barcode technology for Cell Surface Protein. The amplified full-length cDNA was then used to generate the Oxford Nanopore Technology (ONT) long-read sequencing libraries, according to a modified FLT-seq protocol[8] and using the SQK-LSK114 Ligation Sequencing Kit (Oxford Nanopore Technologies, Oxford, UK). Specifically, each sample was amplified in 5 parallel 50 µl PCR reactions using the following reagents: 10 µl 5x

PrimeSTAR GXL Buffer (Takara Bio Inc., Kasatsu, Japan), 4 µl 2.5 mM dNTP solution (New England Biolabs, Ipswich, USA), 1 µl 10 µM FPSfilA primer (5’-ACTAAAGGCCATTACGGCCTACACGACGCTCTTCCGATCT-3’, Thermo Fisher Scientific, Waltham, USA), 1 µl 10 µM RPSfilBr primer (5’-TTACAGGCCGTAATGGCCAAGCAGTGGTATCAACGCAGAGTA-3’, Thermo Fisher Scientific), 1 µl PrimeSTAR GXL Polymerase (Takara Bio Inc.). The thermocycler was set as follows. Step 1: 98 °C for 30 sec; step 2: 8 cycles of 98 °C for 10 sec, 65 °C for 15 sec, 68 °C for 8 min; step 3: 68 °C for 10 min; step 4: 10 °C hold. The pooled PCR product was cleaned up using 0.8x AMPure XP beads (Beckman Coulter Inc., Brea, USA). Specifically, 200 µl AMPure XP beads were mixed with the 250 µl PCR product and incubated at room temperature for 10 minutes. Placing the tube on a magnetic rack cleared the mixture and the supernatant was discarded. The beads were washed with 200 µl 80 % ethanol twice. The beads were airdried for 1 minute after the second ethanol wash. The beads were then resuspended in 51 µl Buffer EB (Qiagen N.V. Hilden, Germany) with the tube off the magnet. Following 5 minutes incubation at room temperature, the tube was placed on the magnet and 50 µl of the clear supernatant was transferred to a new tube. The cleaned-up amplified cDNA was quantified using the Qubit dsDNA HS assay (Thermo Fisher Scientific) and the Fragment Analyzer HS NGS Fragment Kit (Agilent Technologies Inc., Santa Clara, USA).

200 ng of the cleaned-up amplified cDNA was used as an input for library preparation according to SQK-LSK114 Ligation Sequencing Kit (Oxford Nanopore Technologies, Oxford, UK) with modifications. The revised steps of document genomic-dna-by-ligation-sqk-lsk114-GDE_9161_v114_revU_29Jun2022-promethion are highlighted below, in which the values used in the present study are specified in square brackets. DNA repair and end-

prep, step 6: “Using a thermal cycler, incubate at 20°C for 5 [20] minutes and 65°C for 5 [20] minutes”. DNA repair and end-prep, step 10: “Incubate on a Hula mixer (rotator mixer) for 5 [15] minutes at room temperature”. Adapter ligation and clean-up, step 9: “Add 40 [50] µl of resuspended AMPure XP Beads (AXP) to the reaction and mix by flicking the tube”.

We sequenced 50 fmol of library per sample using the Promethion R10.4.1 flowcell, according to SQK-LSK114 Ligation Sequencing Kit (Oxford Nanopore Technologies, Oxford, UK).

The current implementation of Cell Ranger is incompatible with the processing of Nanopore data. The Nanopore data was therefore processed using the wf-single-cell (v.1.0.3) pipeline from Oxford Nanopore Technologies (https://github.com/epi2me-labs/wf-single-cell). The GRCh38 reference package distributed by 10X Genomics (https://www.10xgenomics.com/), "refdata-gex-GRCh38-2024-A", was used.

The filtered gene-level count tables were imported to R (version 4.3.3) and analyzed with Seurat (version 4.3.0.1). The analysis was performed using RStudio (version 2024.12.0+467). We only included the cell barcodes that were identified as containing high quality cells according to the short-read-based analysis. The data was processed as in the bioinformatic analysis section above, starting from gene expression normalization.

The Ensemble ID ENSG00000286122 corresponds *LINC02964* in the Nanopore dataset and *AC016074.2* in the Illumina dataset. The gene name *LINC02964* is used refer to ENSG00000286122 in both datasets.

Due to the differences in the processing pipelines between the Illumina and Nanopore data, we note that the observed differences in gene expression are not necessarily attributed to the capture of longer reads.

*Data access*

Processed sequencing data is accessible through the Gene Expression Omnibus accession number GSE287976. Links to the online web resources are available through the accession number, and at http://basoscrna1.dahlinlab.cmm.se, http://basoscrna2.dahlinlab.cmm.se, http://basoscrna3.dahlinlab.cmm.se, and http://basoscrna4.dahlinlab.cmm.se, in which the numbers indicate each respective figure. Access to raw sequencing data is regulated by the Swedish Ethical Review Authority, Swedish legislation, and General Data Protection Regulation. Access to such data is provided by the corresponding author, given that all legal requirements are fulfilled.

### Supplementary Tables

**Table E1: Total number of sequencing reads per sample**

| Dataset (donor) | Gene Expression Total number of Illumina reads | Gene Expression Total number of Nanopore reads | Surface Protein Total number of reads |
|---|---|---|---|
| Fig 1 (DonorA) | 422 441 331 | NA | 73 536 058 |
| Fig 2 (DonorB) | 392 640 438 | NA | 80 885 932 |
| Fig 3 and 4 (DonorC) | 724 347 603 | 95 369 616 | 276 126 031 |
| Fig 3 and 4 (DonorD rep1) | 786 665 522 | 71 790 574 | 260 983 738 |
| Fig 3 (DonorD rep2) | 750 427 247 | NA | 351 590 581 |

NA, not applicable

**Table E2: Cutoffs used for the quality control filtering of each dataset**

| Dataset (donor) | Features | Percent mitochondria | CD45 reads | PE+ | PE- |
|---|---|---|---|---|---|
| Fig 1 (DonorA) | 500 ≤ Features ≤ 6500 | ≤ 8 | ≥ $10^{2.5}$ | ≥ $10^{2.9}$ | ≤ $10^{2.2}$ |
| Fig 2 (DonorB) | 500 ≤ Features ≤ 3500 | ≤ 6 | ≥ $10^{2.5}$ | ≥ $10^{2.3}$ | ≤ $10^{1.5}$ |
| Fig 3 (DonorC) | 500 ≤ Features ≤ 3500 | ≤ 8 | ≥ $10^{1.5}$ | NA | NA |
| Fig 3 (DonorD_rep1) | 500 ≤ Features ≤ 3500 | ≤ 8 | ≥ $10^{1.5}$ | NA | NA |
| Fig 3 (DonorD_rep2) | 500 ≤ Features ≤ 3500 | ≤ 8 | ≥ $10^{1.5}$ | NA | NA |

NA, not applicable.

**Online repository Figure legends**

**Fig E1.** Detection of basophils among low-density leukocytes. **(A)** Distribution of the CD45-derived reads on $\log_{10}(x+1)$ scale, based on the unfiltered Cell Ranger output. The line corresponds to the cutoff set to distinguish cell-containing droplets and empty droplets. **(B)** Distribution of the isotype control-derived reads on $\log_{10}(x+1)$ scale, based on the unfiltered Cell Ranger output. **(C)** Violin plots showing the number of unique molecular identifiers (UMIs) after quality control filtering.

**Fig E2.** The c-Kit$^{+}$ basophils do not constitute a transcriptionally distinct basophil population. **(A)** Flow cytometry analysis showing the proportion of barcoded cells prior to CITE-seq. The plots show live cells. **(B)** Analysis of CITE-seq data showing the proportion of barcoded cells. The histogram shows the distribution of PE-oligonucleotide-derived sequencing reads after the quality control filtering steps of the pipeline. **(C)** Bar plots showing the fraction of c-Kit$^{+}$ basophils in the two transcriptionally distinct basophil populations.

**Fig E3.** The two transcriptionally distinct populations are independently identified in the three replicates. **(A)** Flow cytometry analysis of leukocytes. The analysis was performed omitting or following Ficoll gradient centrifugation. The gating strategy of a representative experiment is shown on the left (plots show live singlets) and the quantification of basophils and eosinophils of 5 donors from 5 independent experiments is shown on the right. **(B)** Flow cytometry analysis of the cells prior to CITE-seq. **(C)** UMAP visualization colored based on the independent CITE-seq libraries. **(D)** Cells colored accorded to the cell cycle phase. **(E)** UMAP visualizations of the 3 individually prepared transcriptomic libraries processed independently without any data integration. The colors represent Leiden clusters calculated for each dataset separately at resolution 0.1. **(F)** UMAP visualizations of the normalized gene expression of the 3 individually prepared libraries.

**Fig E4.** Immunoprofiling of the circulating human basophils. **(A)** Ridge plots showing CLR-normalized cell surface epitope expression (range 0 to 7, with 2 units between ticks) for the cells annotated as Basophils 1 or Basophils 2 (pink) compared with the other cell types (grey). The cell assignment was performed based on the harmonized dataset and the clustering shown in Figure 3B. The values below the vertical dashed line were considered background, based on the isotype controls (first row). The plots were generated by using the function RidgePlot with default parameters. **(B)** Ridge plots showing the normalized protein expression of the four proteins that showed a bimodal expression in basophils. The plots show the basophils (Basophils 1 and 2 aggregated) of each donor. **(C)** UMAP visualization showing the CLR-normalized protein expression of CD352.

**Fig E5.** Identification of transcriptional heterogeneity by long-read sequencing. **(A)** Dot plots of the correlation between the unique molecular identifiers (UMIs) per cell in the short- and long- read sequencing. Cor indicates Spearman correlation between the two sequencing technologies. **(B)** UMAP visualization of the long-read dataset colored by donor. **(C)** UMAP visualization of the dataset of Fig 3, showing only the cells present in the long-read sequencing dataset. The cells of the Basophils 1 ONT cluster are colored. **(D)** UMAP visualization of the Nanopore dataset for Donor C processed independently. The colors represent Leiden clusters calculated at resolution 0.2. **(E)** Normalized gene expression of selected genes. **(F)** UMAP visualization of the Nanopore dataset for Donor D processed independently. The colors represent Leiden clusters calculated at resolution 0.2. **(G)** Normalized gene expression of selected genes. **(H)** UMAP visualizations based on the

Nanopore coordinates, showing the normalized gene expression of the short-read-based (Illumina) gene expression.
