## Supplementary figures and images for "Single-cell omics-based characterization of human basophils reveals two transcriptionally distinct populations"

### Fig E1

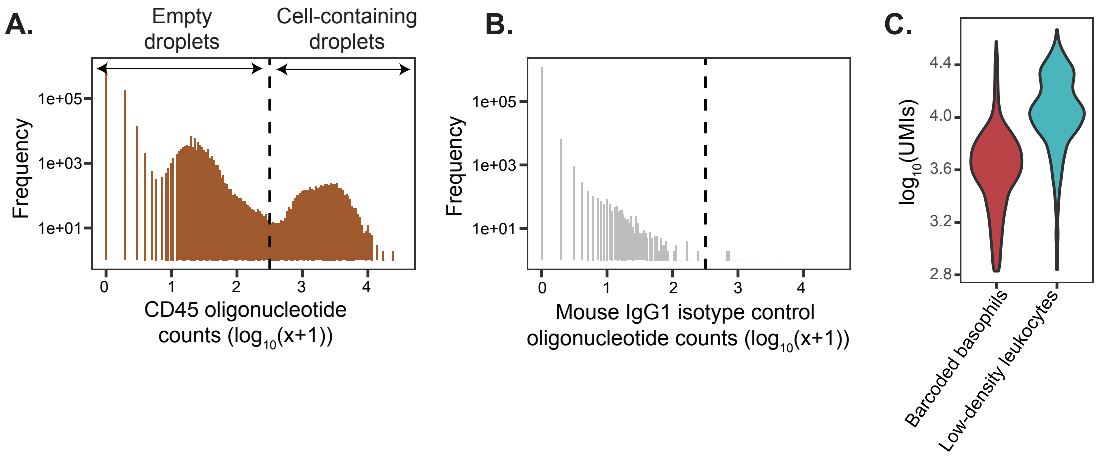

### Fig E2

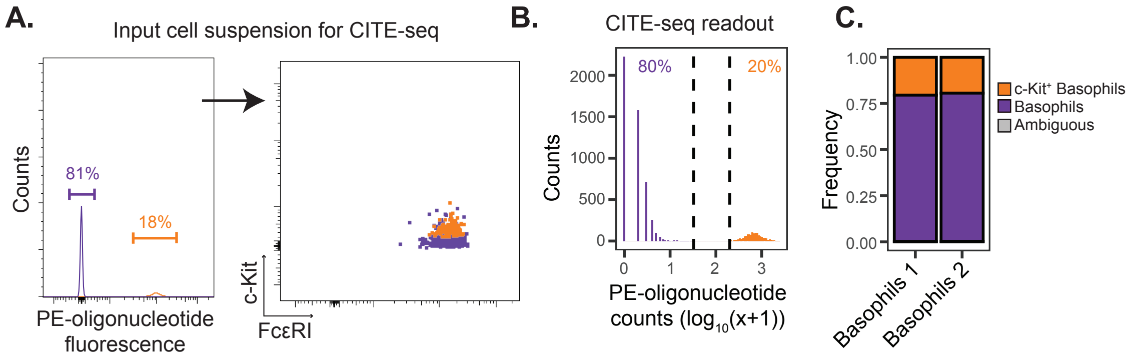

### Fig E3

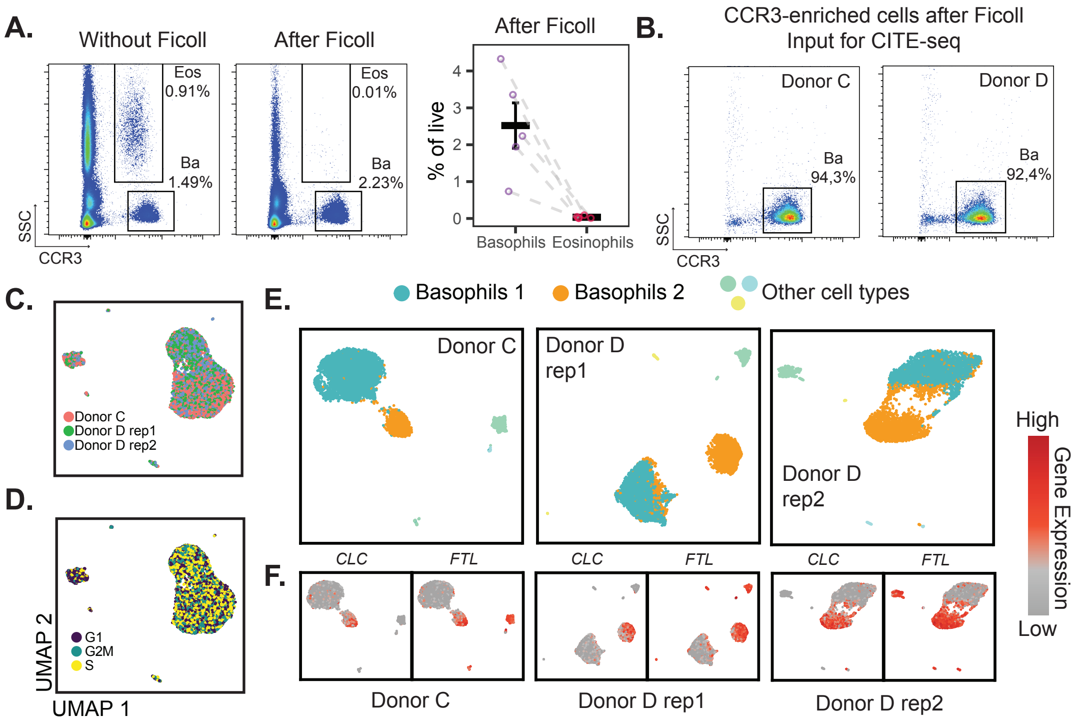

### Fig E4

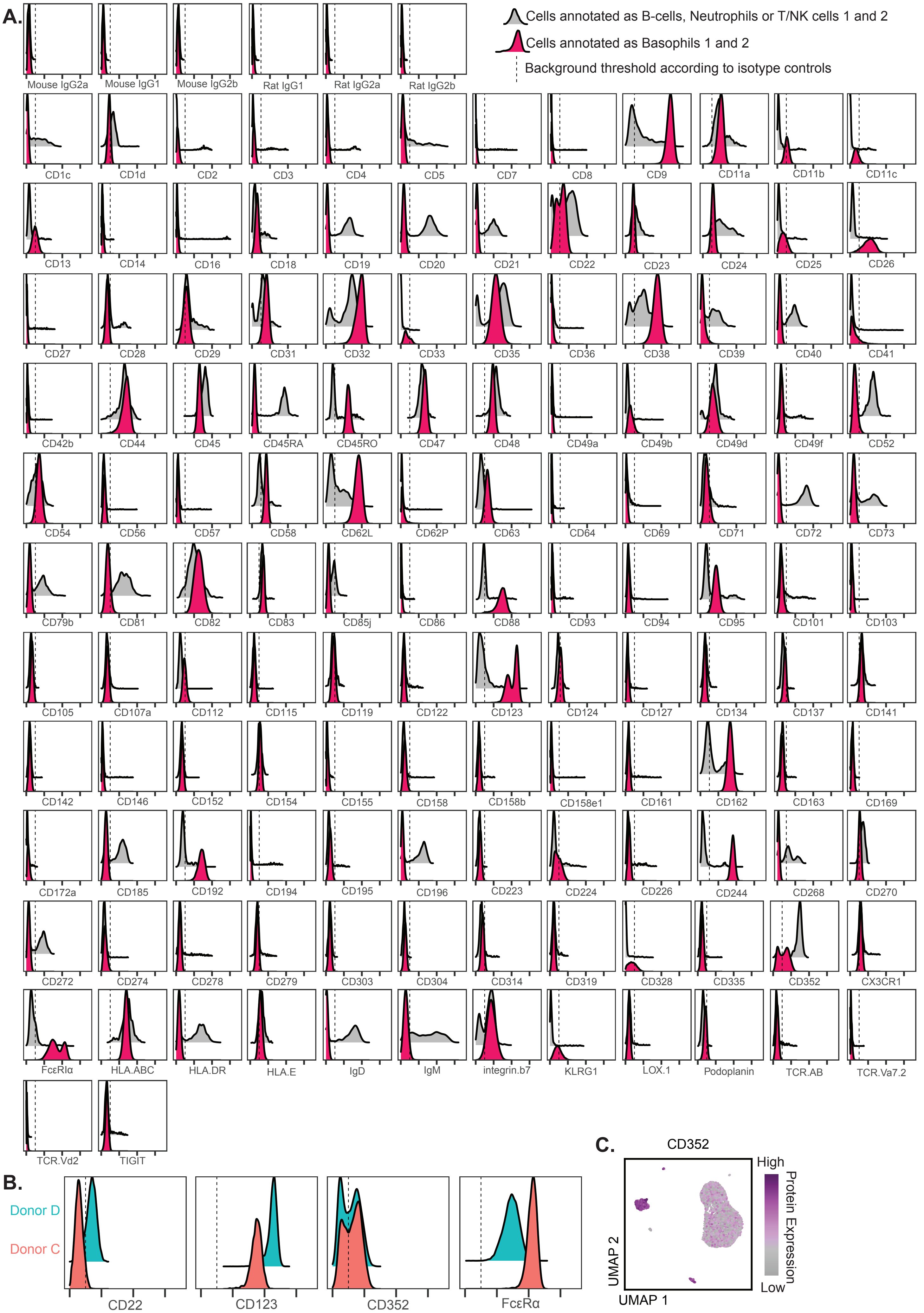

### Fig E5

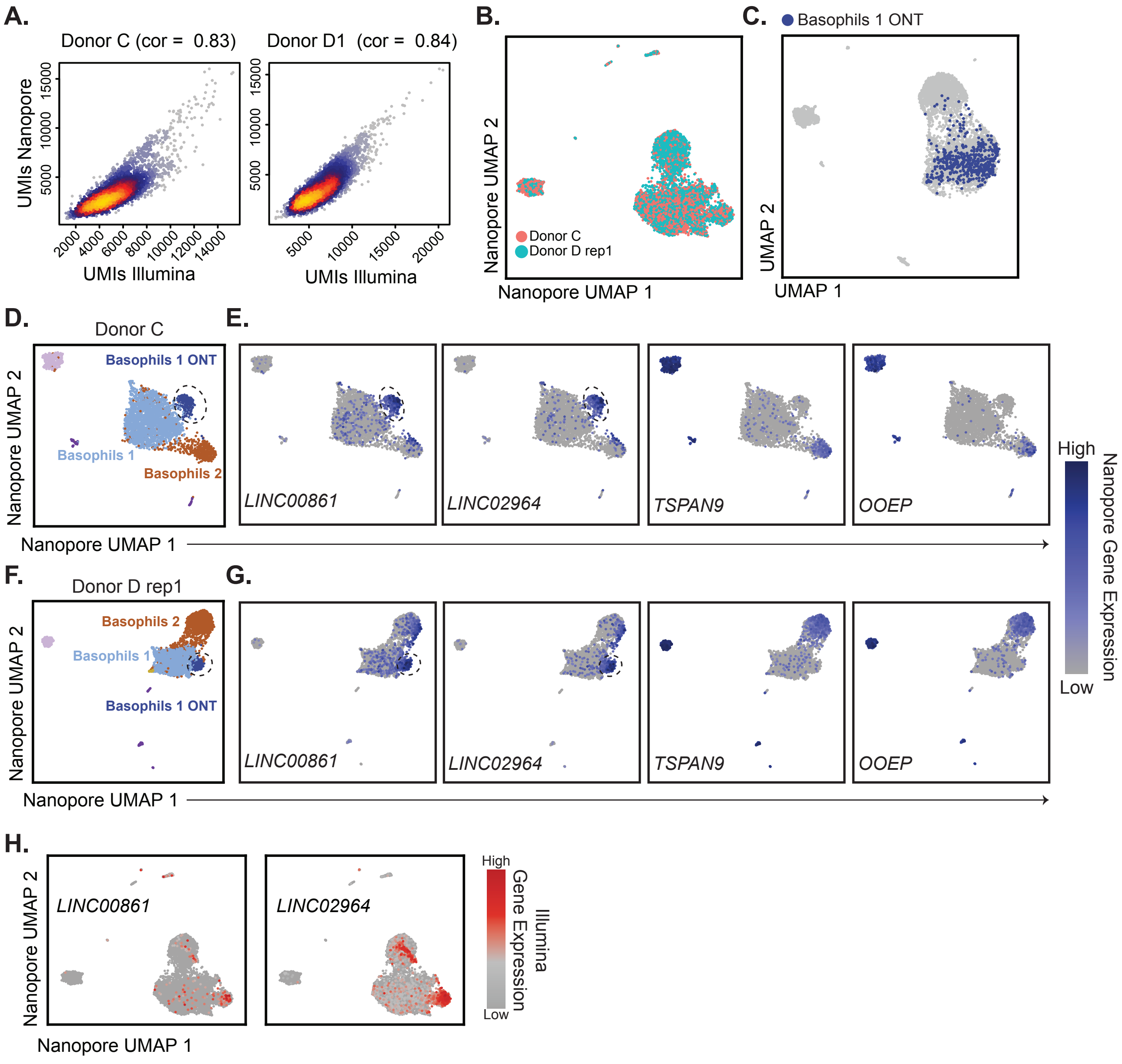
